# Integrated lipidomic and transcriptomic analyses reveal novel endogenous lipid signaling system regulation in skin and plasma during psoriasiform inflammation

**DOI:** 10.64898/2026.05.01.722227

**Authors:** Eva Wisniewski, Wenwen Du, Jenna A Himelstein, Gergo Szanda, Taylor J Woodward, Ken Mackie, Heather B Bradshaw

## Abstract

Psoriasis is a chronic inflammatory skin disease characterized by keratinocyte hyperproliferation and immune dysregulation. Emerging clinical and experimental evidence suggests that endogenous lipid (endolipid) signaling systems, including the endocannabinoid system (ECS), represent a promising therapeutic target to treat psoriasis; however, comprehensive characterization of small-molecule endolipids and related proteins in psoriatic skin and their relationship to systemic changes remains limited.

Here, we used the imiquimod (IMQ)-induced mouse model of psoriasis to perform combined lipidomic and transcriptional profiling of endolipid signaling in both skin and plasma. Targeted lipidomics revealed a striking divergence between tissues: most endolipids increased in inflamed skin but decreased in plasma, including the canonical ECS lipids anandamide and 2-arachidonoylglycerol. In contrast, selected lipid species, including taurine-conjugated metabolites (both *N*-acyl taurines and bile acids), were elevated in both tissues, indicating pathway-specific regulation.

Targeted transcriptional analysis of whole skin showed reduced expression of key endolipid biosynthetic enzymes (*Napepld, Dagla, Daglb*) and the cannabinoid receptor *Cnr1*, while *Cnr2* and ECS-related metabolic enzymes remained unchanged. Additional alterations were observed in transcripts involved in related endolipid signaling (*Trpv1, Trpv4, Ppara, Pparg, Gpr55*), bile acid metabolism (*Fxr, Bsep, Fabp4, Fabp5, Cyp27a1, Cyp8b1*), and inflammatory pathways (*Cox-2*). To resolve this apparent discrepancy between lipid levels and gene expression, we performed compartment-specific analyses of epidermal and dermal layers. These revealed a predominantly suppressive epidermal response across multiple ECS-related proteins, contrasted by a more variable dermal profile with selective preservation or upregulation, particularly of Cnr2.

Together, these findings demonstrate that psoriasiform inflammation is associated with compartment-specific remodeling of endolipid signaling across skin and systemic compartments, underscoring the functional heterogeneity of epidermal and dermal layers. This dataset provides novel insights into the dysregulation of endolipid signaling systems in psoriasis and provides a foundation for the development of spatially informed, lipid-based therapeutic strategies.

## Introduction

Psoriasis is a chronic, immune-mediated skin disease affecting approximately 2-3% of the global population [1]. Emerging clinical evidence suggests that cannabinoids may have therapeutic potential in this disease: topical cannabidiol (CBD) has been reported to improve psoriatic lesions and reduce symptoms in human studies, suggesting a therapeutic role for the endocannabinoid (ECS) system [2–5]. These observations highlight endogenous lipid (endolipid) signaling pathways as promising targets in psoriasis.

Clinically, psoriasis presents as erythematous, scaly plaques driven by keratinocyte hyperproliferation and immune cell infiltration [6, 7]. At the molecular level, this process is sustained by inflammatory cytokines including IL-17, IL-22, IL-23, and TNF-α, which mediate pathogenic immune-keratinocyte crosstalk [8, 9]. The disease arises from a complex interplay of genetic and environmental factors, such as infection, stress, or mechanical injury [6, 7]. Beyond cutaneous manifestations, psoriasis is associated with systemic comorbidities such as psoriatic arthritis, cardiovascular disease, metabolic syndrome, and depression, resulting in substantial physical, psychosocial, and economic burden [10–16]. Although advances in targeted therapies have improved disease management, long-term treatment remains challenging, underscoring the need for safer and more accessible strategies, especially topical treatments. A deeper understanding of disease mechanisms is therefore essential to identify new therapeutic targets. In this context, alterations in endolipid metabolism have emerged as a hallmark of psoriatic skin. Previous lipidomic studies have largely focused on circulating lipids or immune cells, with limited characterization of endolipids in lesional skin [17–21]. Consequently, integrated analysis of these endolipid mediators across both skin and systemic compartments remains lacking [22].

The cutaneous ECS is an endolipid signaling network expressed in multiple skin cell types [23–26] that contributes to skin homeostasis. The canonical ECS comprises the cannabinoid receptors CB1 and CB2, endogenous ECS ligands, including anandamide (AEA) and 2-arachidonoylglycerol (2-AG), and the enzymes responsible for their synthesis and degradation [27, 28]. Beyond these canonical ligands, a broader class of structurally related bioactive lipids - collectively referred to here as endolipids - share metabolic pathways and functional overlap with ECS components. These include more than 100 AEA and 2-AG congeners and related lipid mediators that are regulated by ECS-associated enzymes and contribute to lipid signaling networks in the skin. ECS-related signaling regulates keratinocyte proliferation and differentiation, inflammatory responses, lipid metabolism, and sensory processes such as pain and itch [29–32].

Dysregulation of ECS components has been reported in psoriasis, including altered expression of ECS-related genes [33], altered receptor expression in lesional skin [34], and changes in circulating endolipid levels [19, 35]. In preclinical models, endolipids, as well as synthetic and phytocannabinoids, exert anti-inflammatory, anti-proliferative, and anti-pruritic (anti-itch) effects in the skin [36–43], and experimental studies indicate a protective role for CB2 signaling in psoriasiform inflammation [44–46].

Endolipid signaling is integrated into the broader lipid mediator networks that likely play regulatory roles in disease progression. Bile acids have emerged as signaling molecules that can regulate inflammation and metabolism beyond their classical digestive roles. Conjugated bile acids can modulate N-acyl phosphatidylethanolamine-specific phospholipase D (NAPE-PLD), linking bile acid signaling to endolipid biosynthesis [47, 48]. In addition, bile acids regulate nuclear receptors such as FXR and PPARα and influence pathways involved in inflammation and itch [49]. We recently showed that endolipids are significantly dysregulated in adipose tissue in an FXR KO model that is exacerbated by obesity [50]. Modulation of bile acid signaling has also been reported to ameliorate psoriasiform dermatitis in experimental models [51, 52]. These observations suggest that inflammatory skin disease may involve coordinated remodeling of interconnected endolipid signaling pathways rather than isolated changes within a single system.

Topical application of imiquimod (IMQ) is a widely used murine model of psoriasis [53–55]. IMQ acts as a TLR7/8 agonist and induces erythema, scaling, keratinocyte hyperproliferation, and immune cell infiltration, accompanied by activation of the IL-23/IL-17 axis [56, 57]. Using this model, we evaluated the effects of psoriasiform inflammation on endolipid signaling systems in skin, including selective analyses of the epidermis and dermis, as well as in plasma, to better understand local and systemic endolipid mediator dynamics and their potential relevance to psoriasis.

## Materials and Methods

### Subjects

Ten-week-old male C57BL/6J mice were purchased from The Jackson Laboratory (*Bar Harbor, ME, USA*) and acclimatized for one week prior to experimentation. Mice were housed under a 14/10-hour light/dark cycle at room temperature (22 ± 1 °C) with *ad libitum* access to food and water. All experimental procedures were approved by the Institutional Animal Care and Use Committee (IACUC) of Indiana University (protocol number 24-037).

### Imiquimod psoriasis model procedure

Psoriasis-like skin inflammation was induced using topical imiquimod (IMQ) as previously described [56]. In brief, depilation of a 2 × 3 cm area on the dorsal back skin was achieved using Nair, then a daily dose of 62.5 mg of 5% IMQ cream (*Taro Pharmaceuticals, Israel*) was applied to the back and right ear for six consecutive days. In the control group, the dorsal skin was depilated but left untreated (to avoid potential vehicle-induced alterations in baseline skin lipid composition). Mice were evaluated daily for clinical signs of inflammation under isoflurane anesthesia. Back skin was scored for erythema (redness), scaling, and thickness using a semiquantitative scoring system based on the Psoriasis Area and Severity Index (PASI): 0 = none, 1 = slight, 2 = moderate, 3 = marked, and 4 = severe. In addition, body weight and the thickness of back and ear skin were measured daily using a digital micrometer (*Mitutoyo, Japan*).

On day 7 (after six treatments), mice were euthanized by isoflurane anesthesia and cervical dislocation. Two 4 mm-diameter punch biopsies were collected from the back skin and snap-frozen for lipidomic analysis and RNA extraction. Additional back and ear skin samples were fixed in 4% paraformaldehyde (PFA) for 24 hours, then incubated in 30% sucrose for cryoprotection before cryosectioning. Heparinized blood was collected and centrifuged at 3,000 rpm for 10 minutes to isolate plasma, which was stored at −80 °C until further analysis.

For the follow-up experiment assessing mRNA expression in separated epidermal and dermal compartments, the same IMQ treatment protocol was applied to an independent cohort of 10 C57BL/6J mice. On day 7, dorsal skin from treated and control animals was harvested and processed exclusively for epidermis-dermis separation and subsequent mRNA analysis, without collecting additional tissues or plasma, to maximize RNA yield and preserve compartmental integrity.

### Histological analysis

Cryoprotected tissues were embedded in M-1 embedding medium (*Thermo Fisher Scientific, Waltham, MA, USA*) and cryosectioned at 14 *µ*m thickness using a Leica CM1860 cryostat. Sections were stained using a hematoxylin and eosin (H&E) staining kit (*Abcam Limited, Cambridge, UK*), mounted with Permount™ mounting medium (*Fisher Chemical™, Waltham, MA, USA*), and imaged using a Pannoramic MIDI II digital slide scanner (*3DHISTECH, Hungary, EU*).

### Epidermis-dermis separation

After harvesting the back skin, the tissue was cut into smaller pieces and incubated overnight at 4 °C in 5 mg/mL Dispase II (*Sigma-Aldrich/Merck, Burlington, MA*) diluted in EpiLife medium (*Thermo Fisher Scientific, Waltham, MA*) with gentle rotation to facilitate enzymatic separation of epidermal and dermal layers. Following incubation, the epidermis was mechanically separated from the dermis using curved forceps under a stereomicroscope. Epidermal and dermal fractions were immediately transferred into TRI Reagent (*Zymo Research, Irvine, CA, USA*) for RNA stabilization, and RNA was extracted as described below.

### Real-time quantitative PCR

Total RNA was extracted from back skin biopsies and epidermal/dermal layers using the Direct-zol RNA Miniprep Kit (*Zymo Research, Irvine, CA, USA*) according to the manufacturer’s instructions. First-strand cDNA was synthesized from 3 *μ*g of total RNA using the RevertAid H Minus First Strand cDNA Synthesis Kit (*Thermo Fisher Scientific, Waltham, MA, USA*). Quantitative real-time PCR (qRT-PCR) was performed using TaqMan™ single-tube Gene Expression Assay in 96-well plates (*Applied Biosystems, Waltham, MA, USA*) on a QuantStudio™ 7 Flex Real-Time PCR System *(Thermo Fisher Scientific, Waltham, MA, USA*). For the duplex reactions, targets were dye-labeled with FAM™, and GAPDH-VIC™ was used as the internal reference gene, with two technical duplicates per sample.

TaqMan™ assay IDs targeting the mouse genes of interest were as follows:

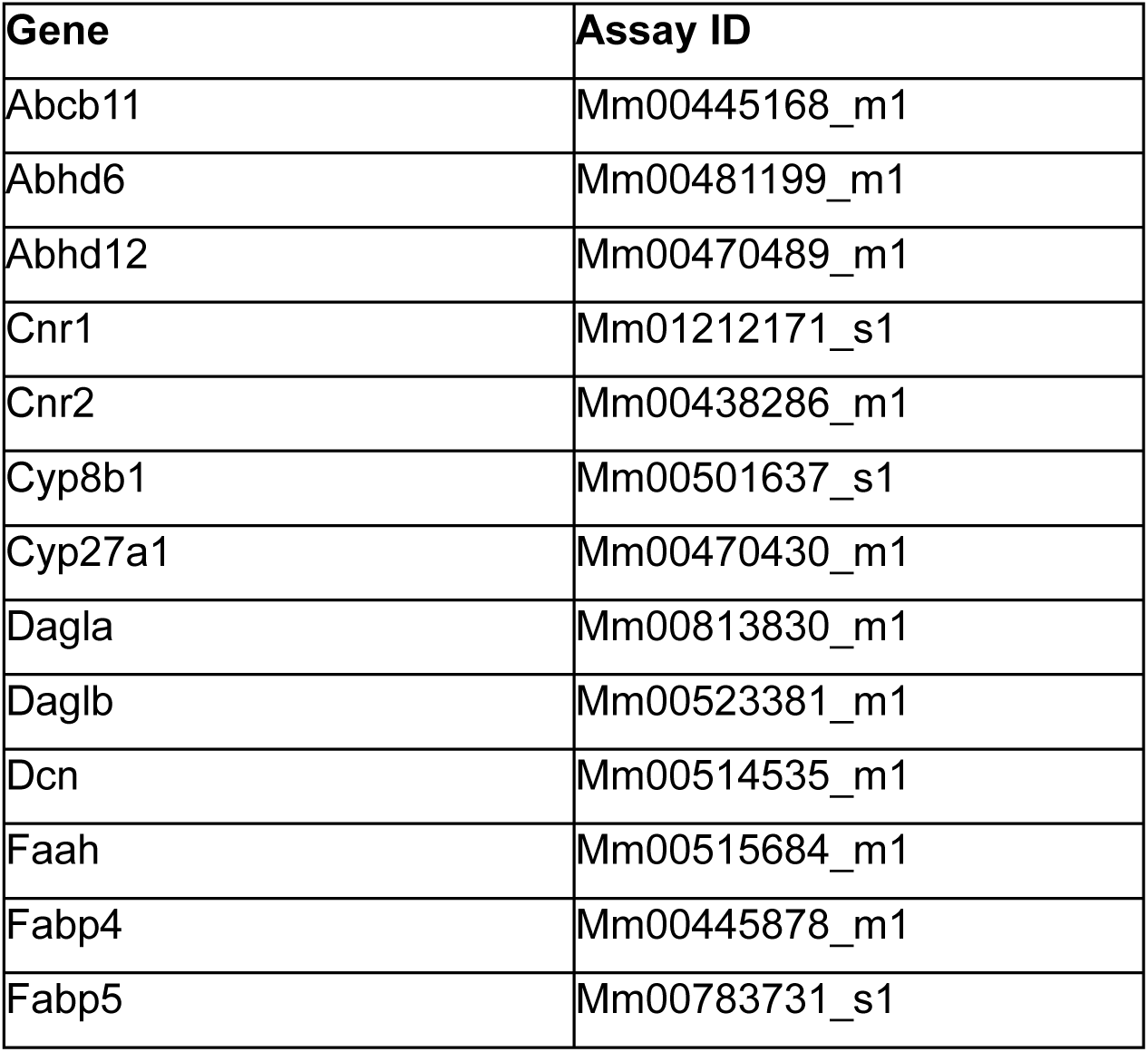
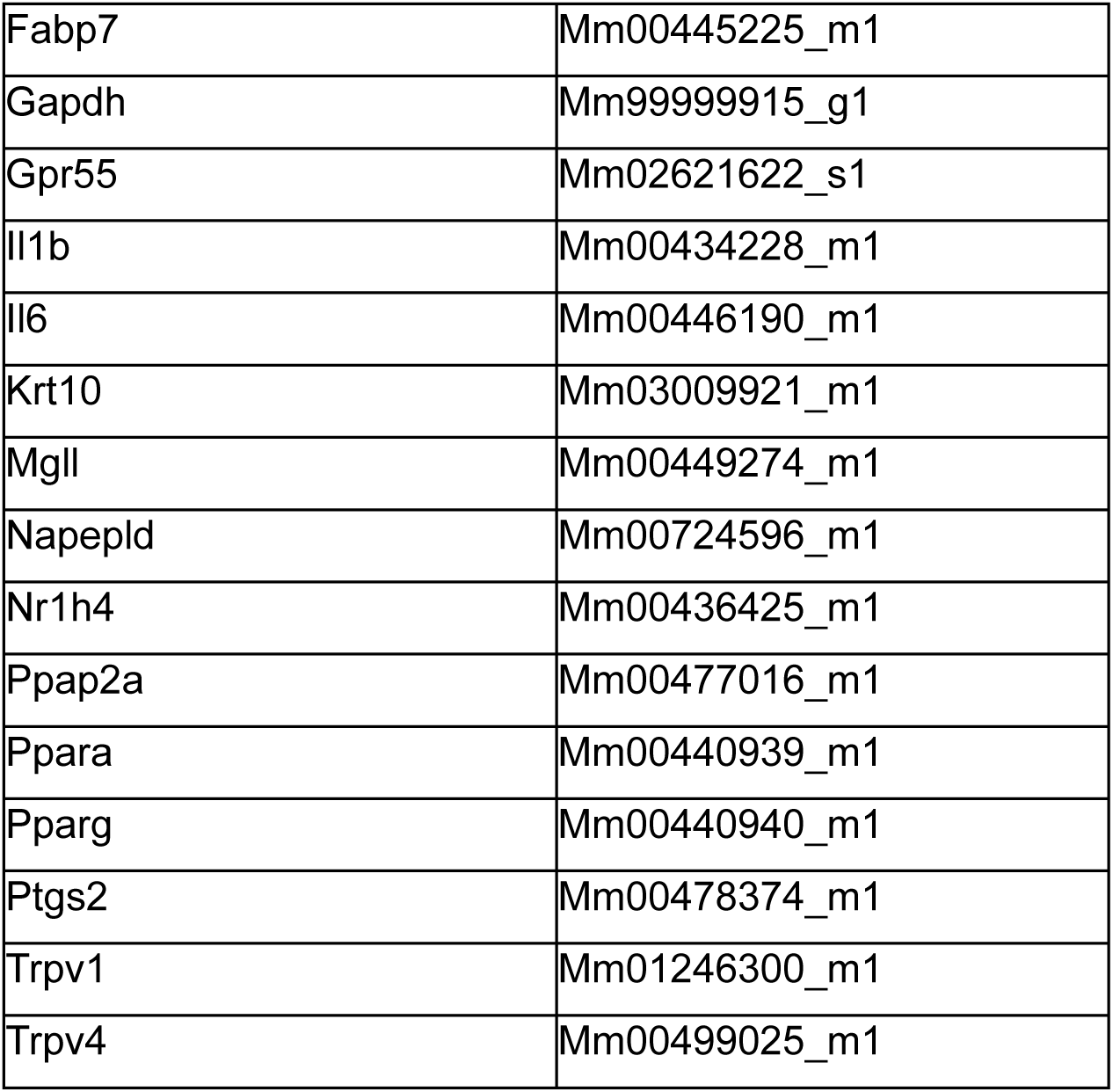

Relative gene expression was calculated using the 2^−ΔΔCt method [58]. Briefly, Ct values for each target gene, measured in technical duplicates, were normalized to the housekeeping gene GAPDH (ΔCt = Ct_target − Ct_GAPDH; measured simultaneously in the same well), and ΔΔCt was calculated relative to the control/calibrator sample or group (ΔΔCt = ΔCt_sample − ΔCt_control average), with expression reported as fold change (2^−ΔΔCt). Outliers were identified using the ROUT method [59]. Statistical analyses and graph generation were performed using GraphPad Prism (*GraphPad Software, San Diego, CA, USA*). Comparisons between two groups were conducted using the non-parametric Mann-Whitney U test. Differences were considered statistically significant at p < 0.05.

### Lipid Extraction and Partial Purification of Plasma and Skin

Lipid extraction and partial purification were performed as previously described [50, 60]. In brief, 50 *μ*L of plasma was added to 2 mL MeOH and 0.10 *μ*L deuterium-labeled anandamide (d8-AEA) and incubated on ice in the dark for 30 minutes. Skin punches were cut into four pieces on an ice-cold dissection plate, then added to 2 mL MeOH and 0.10 *μ*L d8-AEA and sonicated for 60 seconds, then incubated in the dark for 2 hours. Organic supernatants for both tissues were isolated via centrifugation (19,000 G, 20 min, 20 °C), which were then added to 8 mL of HPLC-grade water to form a 20% organic solution. The solution was partially purified with 500mg C-18 solid phase extraction columns (*Agilent, Santa Clara, CA, USA*) with final 1.5 mL elutions of 65, 75, and 100 percent methanol. The methanolic fractions were stored at −80 °C until mass spectrometry (MS) analysis.

### Mass Spectrometry and Lipidomics Analysis

Methanolic elutions were analyzed using HPLC/MS/MS as previously described [50, 60] using the ABI 7500 (*Sciex, Framingham, MA, USA*) coupled to a Shimadzu LC system LC-40DX3 (*Kyoto, Japan*). Standard curves were generated by using purchased standards (*Cayman Chemical, Ann Arbor, MI, USA*) and those made in-house that were validated through NMR and MS analysis as previously described [50, 60, 61]. One hundred and two endolipids were screened using Sciex OS analysis and peak-matching software (*Sciex, Framingham, MA, USA*) coupled to individual chromatographic validation auditing by research staff (*i.e.,* all chromatographic peaks were visualized and verified by an individual prior to data analysis). Statistical significance (p<0.05) and trending (0.05<p<0.1) were differentiated in heatmaps that were generated to visualize changes in the concentration of each lipid analyte between the control and treatment groups as previously described [62]. Students’ T-tests, fold-change, Cohen’s D, and heatmaps were generated using Python (version 3.12.4).

## Results

### Imiquimod-induced psoriasis manifests as thickened, scaly, and erythematous skin in mice

Topical application of imiquimod (IMQ) reliably induced psoriasis-like dermatitis in C57BL/6J mice (Fig. 1). As outlined in the experimental timeline (Fig. 1A), mice received a daily dose of 62.5 mg 5% IMQ cream for six consecutive days, followed by tissue collection on day 7. Macroscopic examination revealed pronounced erythema, scaling, and plaque formation in IMQ-treated animals compared with controls (Fig. 1C, top). Histological analysis of dorsal skin at day 7 further confirmed key features of psoriasiform dermatitis, including epidermal thickening (acanthosis), altered epidermal architecture, and dermal inflammatory cell infiltration (Fig. 1C, bottom). These morphological changes were reflected in quantitative clinical parameters. Daily application of IMQ cream to the depilated back and right ear resulted in progressive increases in ear and back thickness, as well as elevated Psoriasis Area and Severity Index (PASI) scores for erythema, scaling, and thickening (Fig. 1D). Clinical signs became evident by day 3 and continued to increase throughout the treatment period, with erythema reaching its peak on day 6. In contrast, untreated control animals showed no changes in skin thickness or PASI parameters.

**Figure 1.**
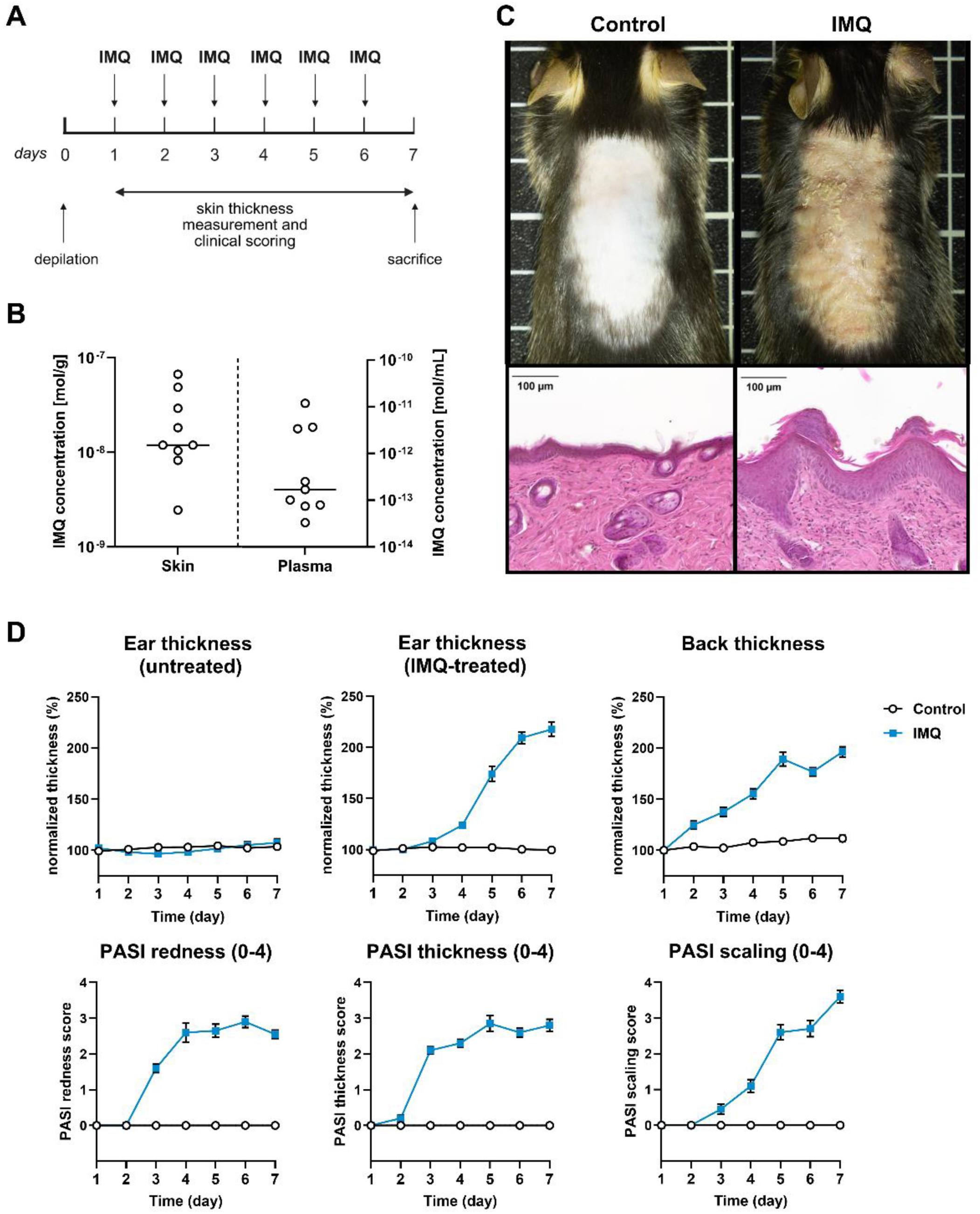
IMQ treatment induces robust psoriasiform dermatitis and confirms local drug exposure. **(A)** Experimental timeline. Mice received 62.5 mg daily topical imiquimod (IMQ) application to the depilated back and right ear for 6 consecutive days. Skin thickness measurements and clinical scoring were performed daily, and tissues were collected on day 7. **(B)** Quantification of IMQ concentrations in skin and plasma 24 h after the final application (day 7) by mass spectrometry. IMQ accumulated in skin, whereas substantially lower levels were detected in plasma. Median is shown. **(C)** Representative dorsal skin images (top) and hematoxylin and eosin (H&E)-stained sections (bottom) from control and IMQ-treated mice at day 7. IMQ-treated skin exhibits erythema, scaling, epidermal thickening, and inflammatory cell infiltration. Scale bars: 100 µm. **(D)** Clinical disease progression during IMQ treatment. Normalized ear thickness (untreated and IMQ-treated ear), back skin thickness, and Psoriasis Area and Severity Index (PASI) parameters (erythema, thickness, and scaling; score 0–4) were assessed daily for 7 days. IMQ treatment resulted in progressive skin thickening and increased PASI scores compared with controls. Data are presented as mean ± SEM (n = 10 per group), thickness was normalized to that measured on day 1 prior to first treatment. Statistical analysis was performed using two-way repeated-measures ANOVA with Sidak’s multiple comparisons test. Significant differences appeared beginning at day 3 and continued throughout the treatment.

To confirm drug exposure, IMQ concentrations were quantified in skin and plasma 24 hours after the final application on day 6. IMQ was readily detectable in both tissues but accumulated at markedly higher (5 orders of magnitude) concentrations in skin than in plasma (Fig. 1B), indicating predominantly local exposure. Notably, plasma IMQ levels did not correlate with those measured in skin across individual animals (Supplementary Fig. S1). Together, these findings confirm robust induction of psoriasiform dermatitis in the cohort used for lipidomic and transcriptional analyses.

### Lipidomics reveals complex tissue-specific changes of endolipids in a model of psoriasiform dermatitis

In the plasma, 83 of the 102 endolipids screened (81%) were detected at analytical limits, and of those, 29% remained unchanged. Of the 71% that significantly changed, 68% significantly decreased upon IMQ treatment. In the skin, 71 of the 102 endolipids screened (70%) were detected at analytical limits, and of those, 27% remained unchanged. Of the 73% that significantly changed, only 8% significantly decreased. Therefore, among IMQ-responsive endolipids, 92% increased in the skin, whereas only 32% increased in plasma. Figure 2 provides examples of individual endolipids in plasma and skin and vehicle control versus IMQ comparisons within tissue types (*e.g.,* fold change, *p*-values). The eCBs AEA and 2-AG are illustrated in 2A and B, respectively, and the AEA congener, oleoyl ethanolamine (OEA: Fig. 2C) fit the profile of decreases in plasma and increases in skin. However, the OEA congener, oleoyl taurine (Fig. 2D), does not fit the pattern in that the levels increase in both skin and plasma. Likewise, the bile acid, cholic acid (CA; Fig. 2E), decreases in both skin and plasma with IMQ treatment, but the taurine-derived CA metabolite, taurocholic acid (Fig. 2F), increases in both tissues in the same way as oleoyl taurine.

**Figure 2.**
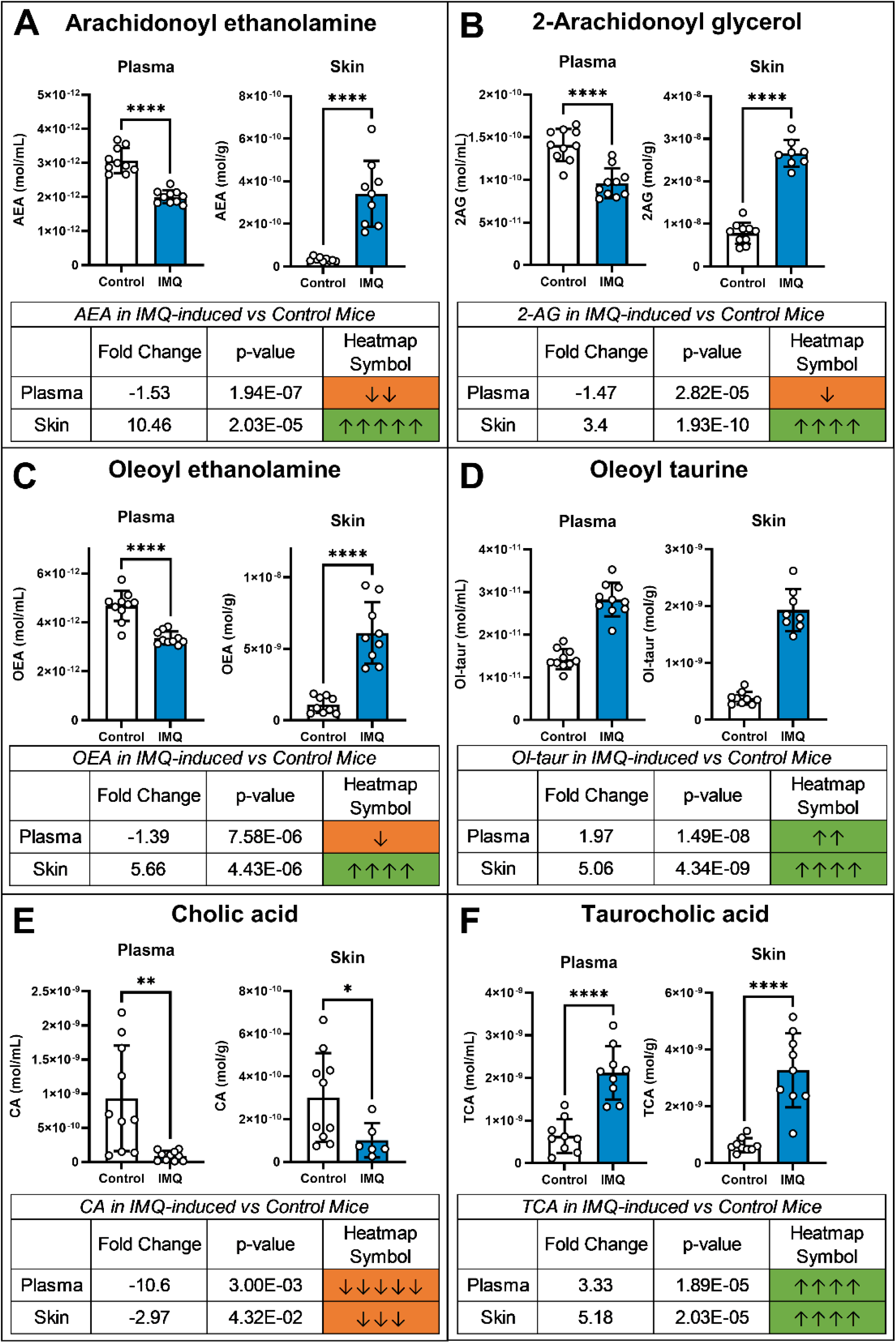
Representative bar graphs of endolipids in plasma and skin with heatmap icons. The bar graphs on the top of the examples are the mean SEM of the endogenous lipid levels in the treatment group (Control in White and IMQ treatment in Blue). **(A)** Arachidonoyl ethanolamine (AEA; Anandamide); **(B)** 2-arachidonoyl glycerol (2-AG); **(C)** Oleoyl ethanolamine (OEA); **(D)** Oleoyl taurine (Ol-taur); **(E)** Cholic acid (CA); **(F)** Taurocholic acid (TCA). Tables underneath the bar graphs list the analytical data of the IMQ vs. control comparisons. *See Methods and Figure 3 for description of Python data analysis for Fold change (denoted by arrows) and t-tests (colors denote significance levels and direction) to generate the Heatmap colors and arrows*.

Figure 3 is a heatmap of all endolipids that were detected at analytical levels in at least one tissue type (*e.g.,* skin, plasma). Endolipids are organized by lipid species type with cholesterol-based bile acids and corticosterone as one cluster, then by the fatty acid group in order of acyl chain length and level of desaturation (*e.g.* palmitic, C16:0; stearic, C18:0; oleic, C18:1; linoleic, C18:2; arachidonic, C20:4; and docosahexaenoic, C22:6). See Methods and Figure 3 for a detailed description of the generation of the heatmaps. Also, see Supplementary Figure S2 for all statistical analytics. These key patterns emerged in the data: 1) Overall, the longer chain, unsaturated fatty acids increased in the skin and decreased in the plasma, whereas the shorter, saturated acyl chain derivatives showed more variability. 2) Stearic acid derivatives were elevated in both skin and plasma with IMQ treatment except for the AEA congener, SEA. 3) The primary exception to the pattern other than acyl chain length was the addition of taurine or glycine as the amino acid conjugate. 4) The CA metabolite, TCA (formed through a conjugation of CA and taurine), is increased at the same rate as CA is decreased in both tissues. 5) Levels of corticosterone and prostaglandin E2 are unaffected by treatment in either tissue.

**Figure 3.**
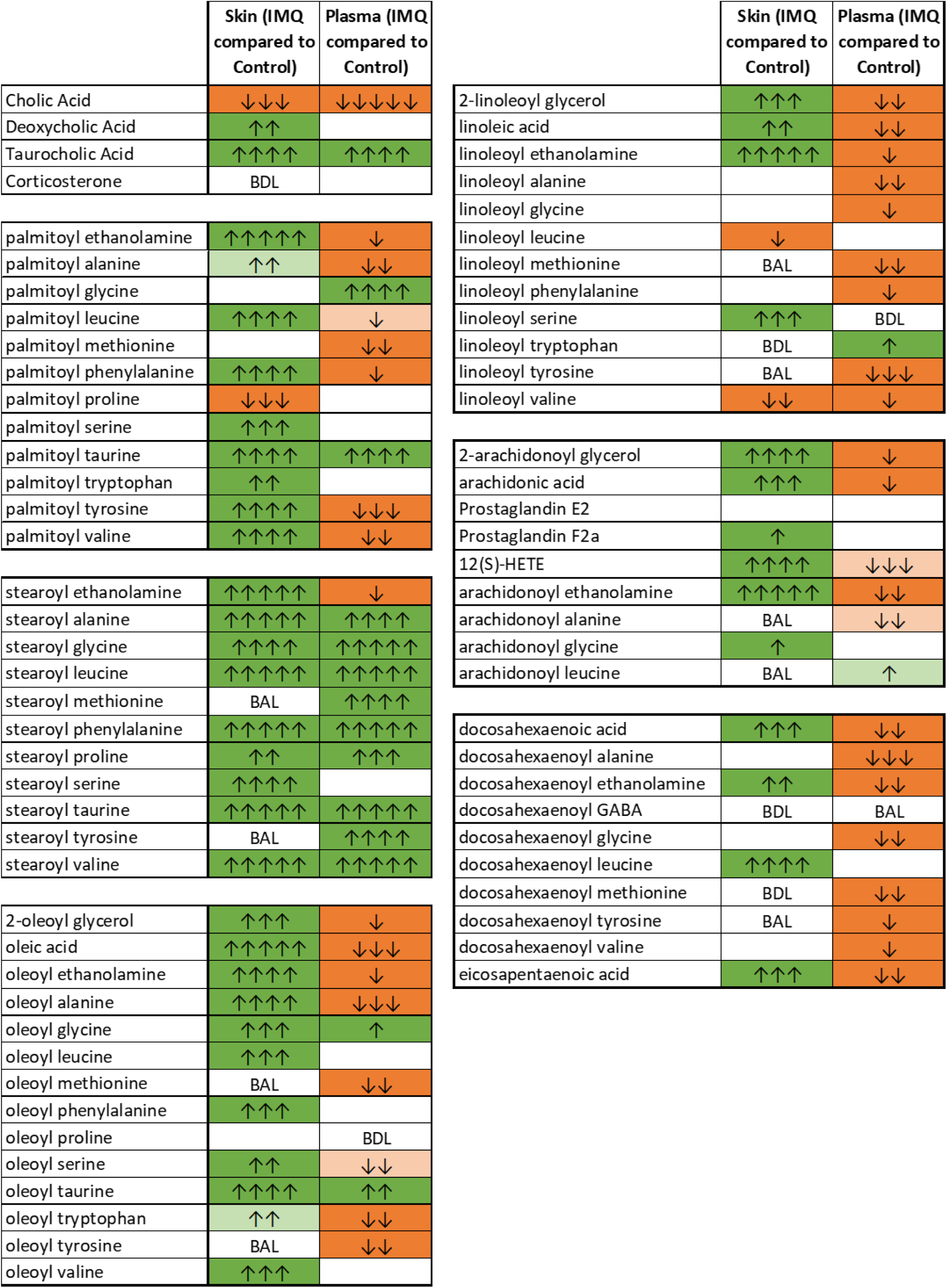
Heatmap of endolipids detected at analytical levels in skin and plasma. Dark green represents significant increases after treatment (p ≤0.05) while light green represents significance levels of p ≤ 0.1-0.05. Dark orange represents significant decreases after treatment (p ≤0.05) while light orange represents significance levels of p≤ 0.1-0.05. Fold change is indicated by the number of arrows, where 1 arrow corresponds to 1–1.49-fold difference, 2 arrows to a 1.5–1.99-fold difference, 3 arrows to a 2–2.99-fold difference, 4 arrows a 3–9.99-fold difference, and 5 arrows a difference of tenfold or more. BAL (below analytical limits) refers to endolipids that were measured in at least one sample but less than 4 making statistical analysis unreliable. BDL (below detectable limits) refers to endolipids that were not detected in any samples. Blank cells indicate that endolipids were present in at least 4 samples in each treatment group being analyzed and that no significant differences were present.

### Psoriasiform inflammation induces coordinated transcriptional changes in endocannabinoid signaling and lipid-associated pathways

To determine whether the alterations in endolipid levels observed in the lipidomic analysis were accompanied by transcriptional changes of endolipid metabolic enzymes, mRNA expression of key ECS components and lipid-related genes were assessed in whole back skin by RT-qPCR. Levels of endocannabinoid-synthesizing enzymes were significantly reduced following IMQ treatment, including the *N*-acyl ethanolamine (NAE)-producing enzyme, *Napepld*, and the 2-acyl glycerol–generating diacylglycerol lipases *Dagla* and *Daglb* (Fig. 4A). In contrast, expression of the primary degradative enzymes fatty-acid amide hydrolase (*Faah*) and monoacylglycerol lipase (*Mgll*) did not differ between IMQ-treated and control animals. Cannabinoid receptor *Cnr1* (CB1) expression was significantly decreased in psoriasiform skin, whereas *Cnr2* (CB2) expression remained unchanged (Fig. 4A).

**Figure 4.**
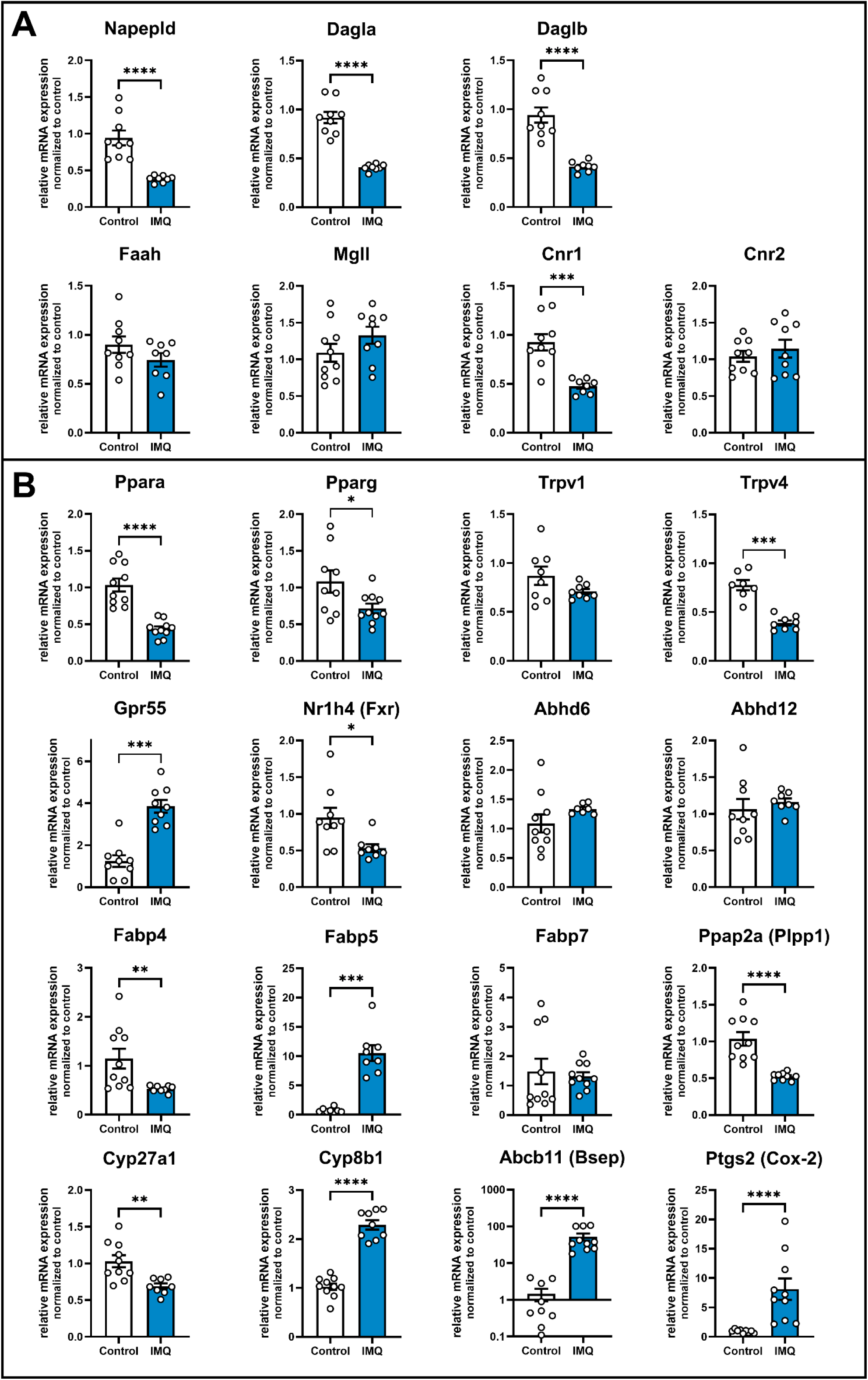
Psoriasiform inflammation drives coordinated transcriptional reprogramming of endocannabinoid and lipid pathways in the skin. Relative mRNA expression was quantified by RT-qPCR in back skin from control and IMQ-treated mice. **(A)** Canonical ECS components, including biosynthetic enzymes (*Napepld, Dagla, Daglb*), degradative enzymes (*Faah, Mgll*), and cannabinoid receptors (*Cnr1, Cnr2*); **(B)** Extended lipid-related pathways, including nuclear receptor signaling (*Ppara, Pparg, Nr1h4/FXR, Ppap2a*), non-classical cannabinoid-related receptors and channels (*Gpr55, Trpv1, Trpv4*), auxiliary endocannabinoid metabolism (*Abhd6, Abhd12*), intracellular lipid transport (*Fabp4, Fabp5, Fabp7*), bile acid synthesis and transport (*Cyp27a1, Cyp8b1, Abcb11*), and inflammatory lipid mediator production (*Ptgs2/COX-2*). Data are shown as individual animals with mean ± SEM (n =10 per group). Statistical significance was assessed using the Mann–Whitney U test. Only statistically significant differences are indicated; absence of annotation denotes non-significant comparisons. *p < 0.05, **p < 0.01, ***p < 0.001, ***p < 0.0001.

To further explore molecular pathways contributing to the endolipid regulation observed in IMQ-treated mice, we reviewed a publicly available RNA-sequencing dataset of IMQ-induced dermatitis across multiple mouse strains [63]. Although the number of animals per group in that dataset was limited, the reported expression patterns for key ECS proteins were largely consistent with the changes observed in our cohort (Fig. 4A). Based on this dataset and their biological relevance, we chose additional endolipid-related genes, involved in nuclear receptor signaling, non-canonical endocannabinoid pathways, lipid transport, bile acid metabolism, and inflammatory lipid mediator production (Fig. 4B).

Within this extended panel, several nuclear receptor-associated pathways were significantly altered. Expression of *Ppara* and *Pparg* was reduced in IMQ-treated skin, as was the bile acid-sensing nuclear receptor *Nr1h4* (FXR), and *Ppap2a* was also significantly decreased. In contrast, expression of the non-classical cannabinoid-related receptor *Gpr55* was markedly increased. Among transient receptor potential (TRP) channels, *Trpv4* expression was significantly reduced, whereas *Trpv1* remained unchanged.

Genes involved in intracellular lipid handling showed differential regulation. *Fabp4* expression decreased, while *Fabp5* was strongly upregulated, and *Fabp7* did not change significantly. Expression of auxiliary endocannabinoid-metabolizing enzymes *Abhd6* and *Abhd12* was not significantly altered.

In pathways related to bile acid metabolism, expression of *Cyp8b1* and the bile salt export transporter *Abcb11* was robustly increased, whereas *Cyp27a1* expression was reduced. Finally, expression of *Ptgs2* (COX-2), a key enzyme in inflammatory lipid mediator synthesis, was markedly elevated in IMQ-treated skin.

Together, these findings show that IMQ-induced psoriasiform inflammation is accompanied by coordinated transcriptional changes extending beyond the canonical ECS, involving lipid transport, nuclear receptor signaling, bile acid metabolism, and inflammatory lipid mediator pathways. Because these analyses were performed in whole-skin samples, they do not resolve compartment-specific effects; however, given the distinct functional roles of epidermal and dermal compartments, differential regulation within these layers is likely and was therefore investigated in subsequent experiments (Fig. 5).

**Figure 5.**
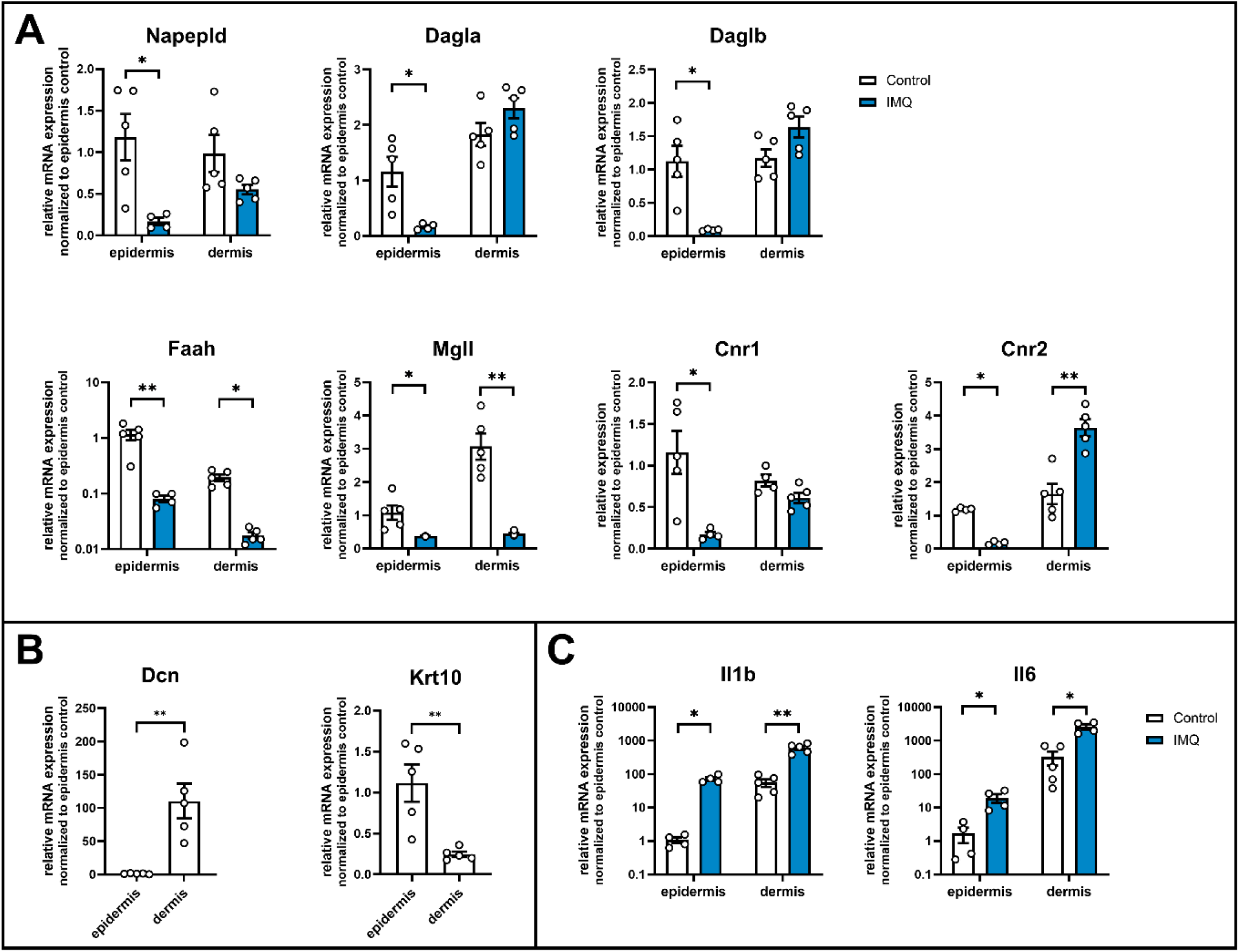
Layer-specific alterations in endocannabinoid system components in epidermis and dermis following IMQ treatment. Relative mRNA expression was quantified by RT-qPCR in enzymatically separated (Dispase II) epidermal and dermal fractions of control and IMQ-treated mice. Data are normalized to control epidermis for visualization. **(A)** Endocannabinoid system components, including biosynthetic enzymes (*Napepld, Dagla, Daglb*), degradative enzymes (*Faah, Mgll*), and cannabinoid receptors (*Cnr1, Cnr2*); **(B)** Validation of tissue separation using epidermal (*Krt10*) and dermal (*Dcn*) marker genes; **(C)** Inflammatory cytokines (*Il1b, Il6*). Data are shown as individual animals with mean ± SEM (n = 4–5 per group). Statistical analysis was performed separately within each skin layer to assess the effect of IMQ treatment (control vs. IMQ) using the Mann–Whitney U test. Comparisons between epidermal and dermal compartments were not performed. Only statistically significant differences are indicated (*p < 0.05, **p < 0.01).

### Layer-specific regulation of endocannabinoid system components in epidermis and dermis

The whole-skin RT-qPCR analysis revealed an unexpected transcriptional pattern, with reduced expression of key endocannabinoid biosynthetic enzymes despite elevated endolipid levels in psoriatic skin (Fig. 4). Given that skin is a heterogeneous tissue composed of distinct cellular compartments with different lipid profiles [64], and that psoriasiform inflammation further alters this composition through keratinocyte hyperproliferation and immune cell infiltration, we next assessed whether epidermal and dermal layers exhibit differential transcriptional responses.

To address this, in an independent cohort, dorsal skin samples were subjected to enzymatic separation of epidermis and dermis before mRNA analysis. Successful separation was confirmed by enrichment of the dermal marker *Dcn* in the dermis and the epidermal marker *Krt10* in the epidermis (Fig. 5B).

Analysis of endocannabinoid biosynthetic enzymes revealed a clear compartment-specific pattern. In the epidermis, expression of *Napepld*, *Dagla*, and *Daglb* was markedly reduced following IMQ treatment. In contrast, dermal expression of these enzymes was maintained or modestly increased, particularly for *Dagla* and *Daglb* (Fig. 5A). A similar pattern was observed for degradative enzymes: *Faah* and *Mgll* expression decreased strongly in the epidermis. In the dermis, Mgll expression was also reduced from higher baseline levels, whereas Faah showed a more modest decrease (Fig. 5A).

Cannabinoid receptor expression further highlighted this compartmentalization. *Cnr1* expression was substantially reduced in the epidermis with comparatively minor changes in the dermis. In contrast, *Cnr2* expression decreased in the epidermis but was significantly increased in the dermal compartment following IMQ treatment (Fig. 5A).

To confirm inflammatory activation within both compartments, expression of *Il1b* and *Il6* were assessed. Both cytokines were robustly induced in epidermal and dermal fractions, with higher expression levels observed in the dermis (Fig. 5C).

Together, these data demonstrate that IMQ-induced inflammation is associated with distinct, layer-specific regulation of ECS-related genes that is not captured in whole-skin analyses.

## Discussion

Understanding how chronic skin inflammatory disorders like psoriasis are driven by alterations in endogenous lipid signaling remains an underexplored avenue for therapeutic development. By combining targeted lipidomics of unique endolipids with transcriptional profiling in the IMQ-induced psoriasis mouse model, we identified novel endolipid profiles in both skin and plasma and showed that endolipid-related enzymes and receptors are differentially regulated across skin compartments. These findings provide a framework for understanding how endolipid signaling is remodeled in psoriatic skin and highlight pathways with potential therapeutic relevance.

### Human psoriasis disease markers and the intersection with the IMQ-mouse model

Human psoriasis is characterized by well-defined morphological and molecular features driven by immune cell-keratinocyte interactions. Histologically, lesions display epidermal hyperplasia (acanthosis) and immune cell infiltration. At the molecular level, disease pathogenesis is largely sustained by activation of the IL-23/Th17 axis, with cytokines such as IL-17A, IL-22, IL-23 and TNF-α promoting keratinocyte proliferation and inflammatory signaling. These processes are further supported by innate immune activation, including Toll-like receptor (TLR) pathways, and are associated with broad transcriptional and metabolic remodeling of the skin [65, 66].

The IMQ-induced mouse model recapitulates many of these hallmark features [55]. Topical IMQ activates TLR7/8 signaling, leading to IL-23/IL-17-driven inflammation, epidermal hyperproliferation, and immune cell infiltration, closely mirroring key histological and immunological aspects of human plaque psoriasis [56]. In addition, transcriptomic analyses have demonstrated substantial overlap between IMQ-induced gene expression signatures and those observed in human psoriatic lesions, particularly in pathways related to cytokine signaling, keratinocyte activation, and immune cell recruitment [53, 57, 63]. These similarities support the translational relevance of the IMQ model for studying psoriasis-associated changes in endolipid signaling.

Within this translational framework, increasing attention has been directed toward lipid-mediated signaling pathways, particularly the cutaneous endocannabinoid system (ECS). ECS-related endolipids and their receptors regulate keratinocyte proliferation, immune responses, and sensory signaling, and growing evidence indicates that modulation of these pathways can attenuate inflammation and restore skin homeostasis [32, 67, 68]. Consistent with this, cannabinoid-based interventions have shown beneficial effects in both preclinical models and early clinical studies of psoriasis [2, 3, 5, 28, 32, 40, 45, 67].

Lipidomic alterations are increasingly recognized as key features of psoriasis at both systemic and tissue levels. Circulating lipid profiles can reflect disease activity, serve as biomarkers or risk factors for comorbidities, and may be used to monitor therapeutic responses [22, 69]. At the same time, the lipid composition of the skin plays a critical role in barrier integrity and redox balance, while specific lipid mediators actively contribute to inflammatory signaling and disease pathogenesis. A deeper understanding of the lipidomic “fingerprint” of psoriasis is therefore critical for improving prevention, diagnosis, and treatment strategies. However, previous human studies have largely focused on either systemic lipid changes (*e.g.*, plasma lipidomics of glycerophospholipids [17], oxylipins/eicosanoids [18], long-chain fatty acids [69], ceramides [21]) or cell- and compartment-specific analyses, such as isolated epidermis [70–72], or immune cells [19, 20]. Comprehensive analyses integrating endocannabinoid-related endolipids across both skin and plasma compartments have not been systematically conducted before.

By performing parallel lipidomic profiling in both skin and plasma, our study addresses this gap and provides an integrated view of local and systemic endolipid changes during psoriasiform inflammation. To complement these measurements and better interpret tissue-level changes, we assessed transcriptional regulation of ECS-related proteins in whole skin and in separated epidermal and dermal layers. This distinction is particularly relevant given the inherent heterogeneity of the skin. The skin comprises multiple metabolically active cell populations – including keratinocytes, fibroblasts, resident and infiltrating immune cells – that contribute differently to lipid signaling networks. Human studies have demonstrated that epidermis and dermis differ substantially in their bioactive lipid composition [64], indicating that whole-skin measurements only represent a composite of functionally distinct compartments. Therefore, a layer-specific analysis provides critical context for interpreting bulk skin data and for understanding how endolipid signaling is differentially regulated across skin compartments during inflammation.

### Lipidomics outcomes with the IMQ model elucidate novel endolipid signaling pathways

The pattern of endolipid regulation in the IMQ model illustrates that the directionality of change in plasma endolipids is the inverse of that for skin endolipids for ∼60% of those detected in the screening library. This finding supports a working hypothesis that specific plasma endolipids, which significantly decrease, are being translocated to the skin because of cellular cues associated with chronic inflammation and that this translocation increases their levels in the skin. The molecular cues for this transfer may involve fatty acid binding proteins that were significantly modulated in the skin here (e.g., FABP4, FABP5). An alternative hypothesis is that the local production and metabolism in the skin is modulated independently to systemic changes. However, here we see decreased NAPE-PLD mRNA in both skin layers, coinciding with *increases* in all *N*-acyl ethanolamines (NAEs) in the skin, consistent with down-regulation and reduced production, likely due to increased local levels. A key difference in the hypothesis that decreases in plasma endolipids are driven by translocation to skin or tissue-specific enzymatic regulations is that stearic acid derivatives in the screen increased more than 3-fold in both tissues, with the single exception of *N*-stearoyl ethanolamine in plasma. The only other endolipids that had increases in both skin and plasma were specific members of the *N*-acyl glycine and *N*-acyl taurine groups. Taken together, this suggests that specific acyl chains (*e.g*., stearic acid) or amino acids (*e.g*., taurine, glycine) have pathway-specific regulation in the context of the IMQ-psoriasis model.

The majority of the endolipids in the lipidomics screen here are small-molecule lipids that are structural analogs to ECS ligands, with many acting as ligands or modulators at GPCRs and TRP channels. Bile acids are a much smaller subset of endolipids in the screen, yet the profound effect on their levels in this model illustrates that multiple endolipid systems are engaged and likely intersect to drive the phenotype. IMQ treatment reduced cholic acid in both plasma and skin, while increasing taurine-conjugated bile acids, including taurocholic acid and tauro(cheno)deoxycholic acid. This shift is notable because bile acids are now recognized as signaling molecules with immunomodulatory functions [73, 74], and bile acid treatment has been reported to improve psoriasiform dermatitis through inhibition of IL-17A production and reduced CCL20-dependent trafficking [51]. Relevant to our data, bile acids can allosterically regulate NAPE-PLD, and glyco- or tauro-dihydroxy conjugates can bind and activate this enzyme [47, 48]. The expression profile of the stearic acid derivatives is like that of the bile acids, being the same in both plasma and skin (either increases or decreases in both), suggesting that the regulatory mechanisms of each of these specific endolipids are more similar than those that show inverse expression.

Two notable endolipids that did not change in either skin or plasma following chronic IMQ treatment were corticosterone and PGE₂. Although PGE₂ is known to regulate keratinocyte proliferation and differentiation and can enhance IL-23/IL-17 signaling [75, 76], we did not observe significant alterations in its levels in either compartment. This is somewhat unexpected given the associated increase in arachidonic acid (AA) and elevated expression of COX-2 in the skin. However, this pattern is consistent with human psoriasis lipidomics, where HETE species, particularly 12-HETE, are more prominently altered than prostaglandins [35, 77]. In line with this, we observed increased 12-HETE levels, which is biologically plausible given its role as a major epidermal AA metabolite with chemotactic and pro-inflammatory functions in the skin [64]. These lipidomic findings suggest that IMQ-induced inflammation does not simply alter endolipid abundance but may also reshape the enzymatic and receptor systems that regulate endolipid signaling. We therefore examined whether these metabolite changes were accompanied by coordinated transcriptional alterations in whole skin and across individual skin compartments.

### Transcriptional outcomes from the IMQ model elucidate novel enzyme and receptor regulatory processes

The whole-skin RT-qPCR data showed a surprisingly consistent transcriptional pattern: expression of the major biosynthetic enzymes NAPE-PLD, DAGLα, and DAGLβ decreased, FAAH and MAGL remained unchanged, and CNR1, but not CNR2, was reduced. This pattern was directionally consistent with the public RNA-seq dataset from IMQ-treated mice [63], supporting its reproducibility. The discrepancy between reduced biosynthetic enzyme expression and increased tissue EC levels indicates that transcriptional changes alone do not predict steady-state endolipid abundance. Potential explanations include post-transcriptional regulation (*e.g*., bile acid-dependent modulation of NAPE-PLD [47]), altered enzymatic activity despite stable mRNA expression, changes in protein half-life, feedback-driven compensatory regulation, or compartment-specific regulation within epidermis and dermis. This interpretation is further supported by our epidermis-versus-dermis analyses from an independent cohort, showing that Dagla and Daglb downregulation is primarily driven by epidermal changes, whereas Napepld decreases in both layers. In this context, the unchanged expression of the major degradative enzymes Faah and Mgll in whole skin similarly does not account for elevated EC levels and should be interpreted with the same caution as biosynthetic enzymes. However, both transcripts were reduced after epidermis/dermis separation, suggesting that bulk skin measurements may mask layer-specific decreases. As with NAPE-PLD, this further supports the notion that enzyme activity and local regulation, rather than transcript abundance alone, are key determinants of endolipid levels. Indeed, several N-acylethanolamines detected in our dataset, including PEA and SEA, share metabolic pathways with anandamide and have been reported to modulate FAAH activity indirectly, for example, through substrate competition or regulation of enzyme expression, raising the possibility that local endolipid composition may further influence degradative capacity [78, 79]. While CB2 has been reported to be increased in both human psoriatic lesions and at least one IMQ-based mouse study [34, 45], our whole-skin analysis did not detect a net increase, consistent with the Swindell RNA-seq dataset [63]. This discrepancy may reflect differences in model conditions and, importantly, the fact that bulk skin measurements average across dynamically changing cell populations. In this context, our finding of decreased epidermal *Cnr2* but increased dermal Cnr2 suggests that spatial redistribution of CB2 receptors, potentially reflecting reduced keratinocyte Cnr2 alongside increased immune cell number/expression, may be more informative and biologically meaningful in psoriasis.

To determine whether this remodeling extended beyond canonical ECS components, we next examined a broader panel of endolipid-responsive receptors, nuclear lipid sensors, auxiliary metabolic enzymes, and genes involved in intracellular lipid handling, bile acid metabolism, and inflammatory lipid signaling.

Beyond CB1 and CB2, our expanded panel suggests that psoriasiform inflammation also remodels non-classical ECS signaling pathways. TRPV4 expression was significantly reduced, whereas TRPV1 remained unchanged. This is notable because several cannabinoid and ECS-related ligands can engage TRPV channels [80–84], and TRPV4 is increasingly recognized as an important regulator of cutaneous barrier, sensory, and inflammatory functions [85]. In psoriasis, reduced TRPV4 expression has been reported in human lesional skin [34], whereas experimental deletion of *Trpv4* in mice attenuates IMQ-induced disease and reduces epidermal thickening and inflammatory cell infiltration, supporting a pathogenic role for this channel in psoriasiform inflammation [86]. In parallel, *Gpr55* was markedly upregulated in our dataset. Although the role of GPR55 in psoriasis remains less well-defined than that of canonical cannabinoid receptors, GPR55 is activated by several cannabinoid-related lipids [87, 88] and has been linked to inflammatory signaling, including NF-κB-related pathways [89]; moreover, increased GPR55 expression has been reported in granulocytes from patients with psoriasis [19].

The changes in nuclear lipid sensors were similarly coherent with a dysregulated anti-inflammatory/metabolic program. Both *Ppara* and *Pparg* expression were reduced, consistent with the reported downregulation of these receptors in human psoriatic lesions [34] and with their established roles in regulating keratinocyte differentiation, lipid metabolism, and inflammatory processes [90, 91]. This is particularly relevant because several endolipids elevated in our study, including AEA, PEA, OEA, and related species, can signal through PPAR pathways [92, 93], and activation of PPARγ has long been associated with anti-proliferative effects in keratinocytes and has therapeutic potential in psoriasis [40, 94–96]. We also observed reduced *Nr1h4* (FXR) expression, which fits with emerging evidence that bile-acid/FXR signaling is protective in psoriasis-like inflammation: experimental enhancement of bile-acid signaling alleviates psoriasiform dermatitis and reduces IL-17-related inflammation [51, 97]. In contrast, ABHD6 and ABHD12, alternative 2-AG hydrolases, were not significantly altered, indicating they are unlikely to be major contributors in this model.

Finally, several changes pointed to broader remodeling of intracellular lipid handling, bile acid metabolism, and inflammatory lipid signaling. *Fabp5* expression was strongly induced, whereas *Fabp4* decreased, consistent with altered intracellular trafficking of bioactive lipids; this is potentially relevant because FABP5 can govern endocannabinoid-related lipid transport and has also been linked to psoriasis severity [98, 99]. *Ppap2a*, which encodes a phosphatase involved in degrading lysophosphatidic acid (LPA), was reduced; although we did not measure LPA directly, this finding is interesting in light of recent work implicating LPA/LPAR5 signaling in IMQ-induced psoriasis-like inflammation [100, 101], raising the possibility that reduced LPA degradation may contribute to inflammatory signaling. The bile acid-related genes were particularly notable: *Cyp8b1* and *Abcb11* were markedly upregulated, whereas *Cyp27a1* was downregulated, indicating reprogramming of bile-acid synthesis and transport. Given that *Abcb11* encodes the bile salt export pump (BSEP) and that our lipidomics showed decreased cholic acid but increased deoxycholic and taurocholic acid species, these transcriptional changes are consistent with a shift in bile-acid handling rather than a passive bystander effect.

### Limitations, conclusions, and future directions

Several limitations of this study should be acknowledged. First, our analyses were performed at a single time point (day 7), reflecting sustained inflammation but not capturing early dynamic changes during disease initiation and progression. Second, although we complemented whole-skin measurements with epidermis/dermis separation, these compartment-specific analyses were conducted in an independent cohort with a smaller sample size and were not directly paired with lipidomic measurements, which may limit quantitative integration across datasets. Third, transcriptional measurements do not necessarily reflect enzymatic activity or lipid flux, and functional studies will be required to determine how these pathways contribute to endolipid regulation in inflamed skin. Finally, while the IMQ model recapitulates many key features of human psoriasis, including IL-23/IL-17-driven inflammation and epidermal hyperproliferation, it remains an acute murine model and may not fully capture the complexity and chronicity of human disease.

Overall, our findings support the concept that the endocannabinoid system and related endolipids represent an important regulatory network in psoriatic skin inflammation. Elevated endolipid levels in psoriatic skin, together with transcriptional alterations in ECS-related proteins, lipid transporters, and bile acid pathways, point toward a coordinated metabolic adaptation within inflamed tissue. These observations are particularly relevant in light of the growing interest in ECS-directed therapies, including synthetic and phytocannabinoids, CB2-selective agonists [102], enzyme inhibitors targeting FAAH or MAGL [103], and multifunctional compounds such as CB2/PPARγ agonists [68, 104]. By integrating lipidomic and transcriptional datasets across skin and plasma, this study defines endolipid-responsive pathways that may represent actionable targets for therapeutic intervention, warranting further validation in human disease.

## Supplementary materials

**Supplementary Figure S1.**
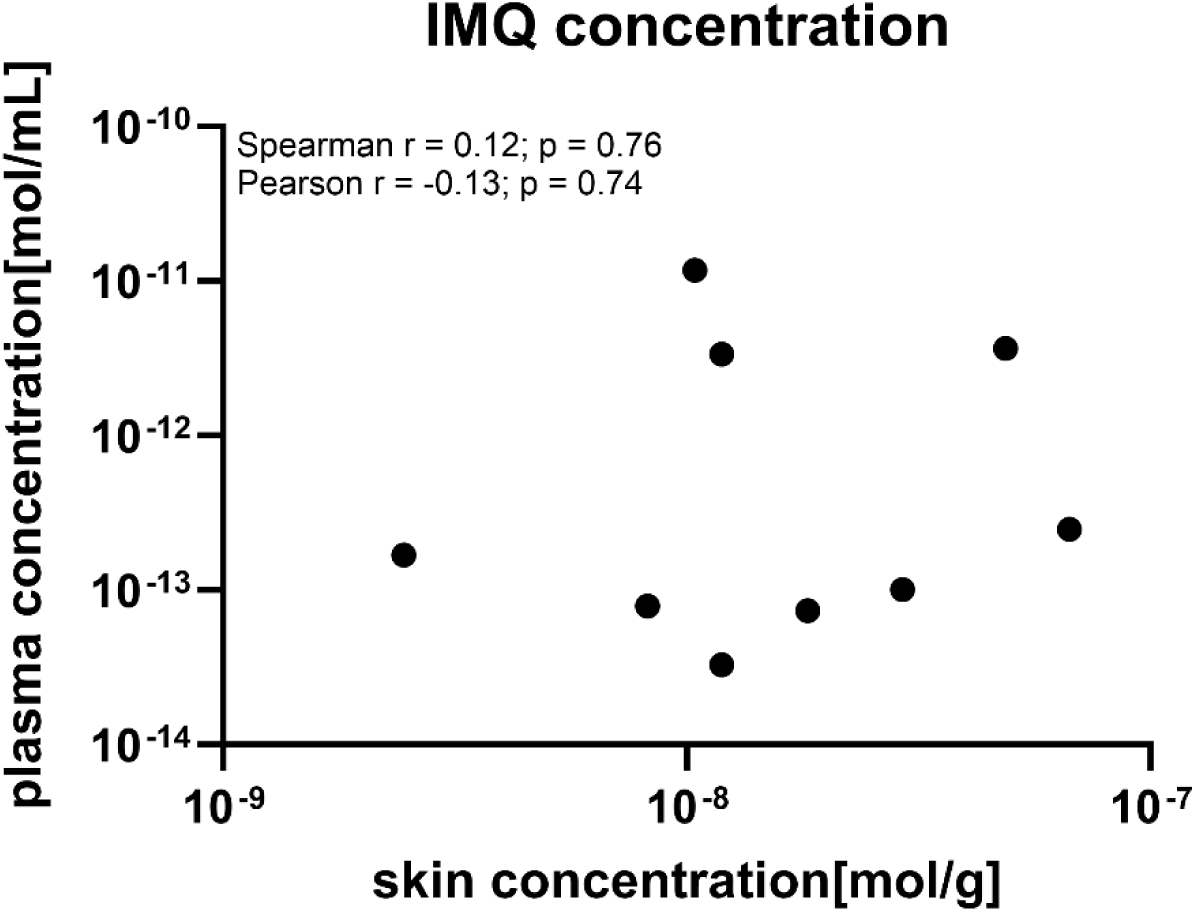
Lack of correlation between skin and plasma imiquimod (IMQ) concentrations. IMQ concentrations were measured in skin and plasma samples collected 24 h after the final topical application. Correlation analysis revealed no significant relationship between skin and plasma IMQ levels across individual animals (n = 9), as assessed by both Pearson (r = −0.13, p = 0.74) and Spearman (r = 0.12, p = 0.76) correlation tests. These data indicate that systemic exposure to IMQ does not directly reflect local skin concentrations under these experimental conditions.

**Supplementary Figure S2.**
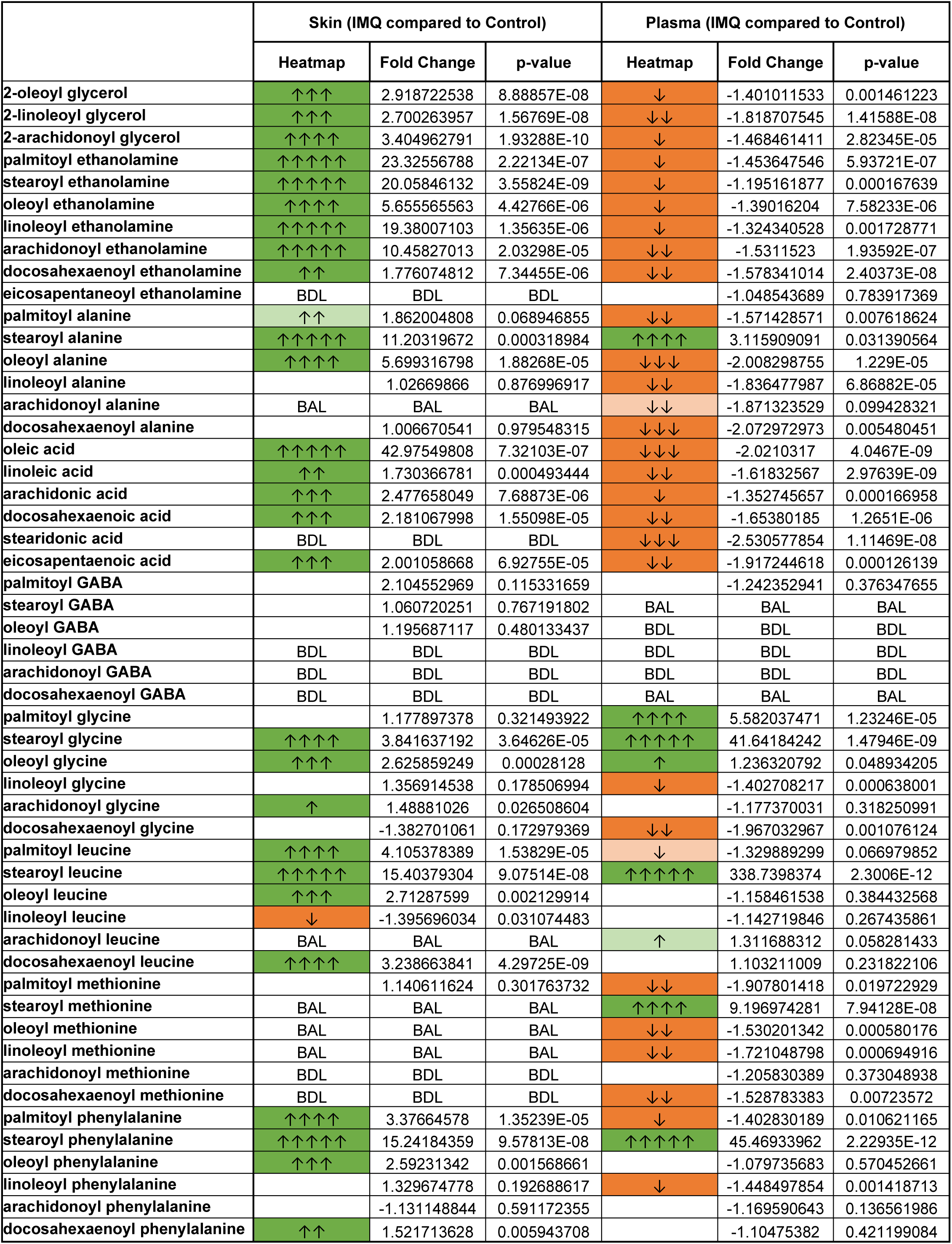
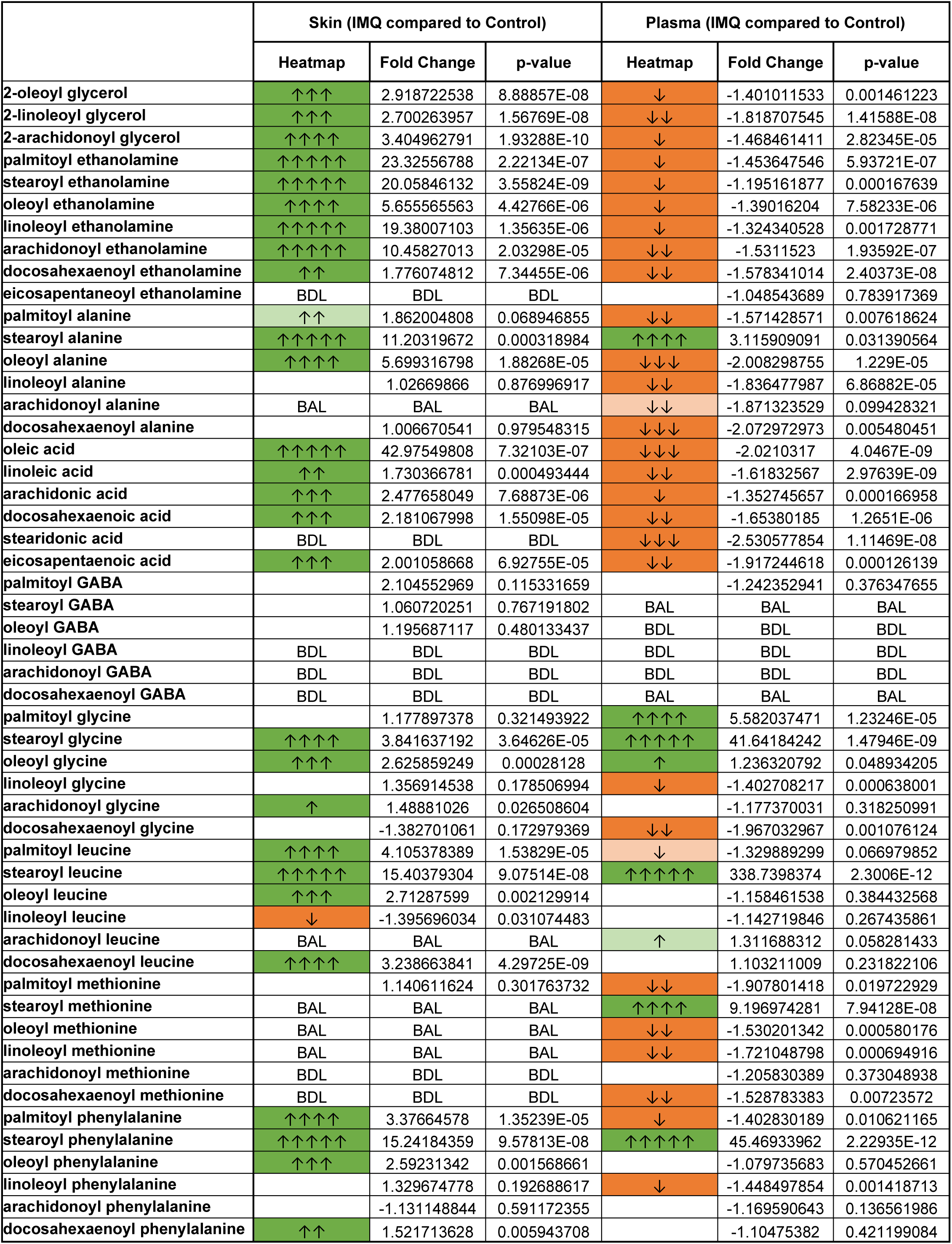
Heatmap, fold change, and statistical analysis of all detected endolipids in skin and plasma following IMQ-induced psoriasiform inflammation. **Supplementary Figure S2. Data analysis of all endolipids scanned in skin and plasma.** The tables list the analytical data of the IMQ versus control comparisons. See Methods for description of Python data analysis for Fold change (denoted by arrows) and T-tests (colors denote significance levels and direction) to generate the Heatmap colors and arrow. Dark green represents significant increases after treatment (p ≤0.05) while light green represents significance levels of p ≤ 0.1-0.05. Dark orange represents significant decreases after treatment (p ≤0.05) while light orange represents significance levels of p≤ 0.1-0.05. Fold change is indicated by the number of arrows, where 1 arrow corresponds to 1–1.49-fold difference, 2 arrows to a 1.5–1.99-fold difference, 3 arrows to a 2–2.99-fold difference, 4 arrows a 3–9.99-fold difference, and 5 arrows a difference of tenfold or more. BAL (below analytical limits) refers to endolipids that were measured in at least one sample but less than 4 making statistical analysis unreliable. BDL (below detectable limits) refers to endolipids that were not detected in any samples. Blank cells indicate that endolipids were present in at least 4 samples in each treatment group being analyzed and that no significant differences were present.

