## Supplemental Figures for "Integrated lipidomic and transcriptomic analyses reveal novel endogenous lipid signaling system regulation in skin and plasma during psoriasiform inflammation"

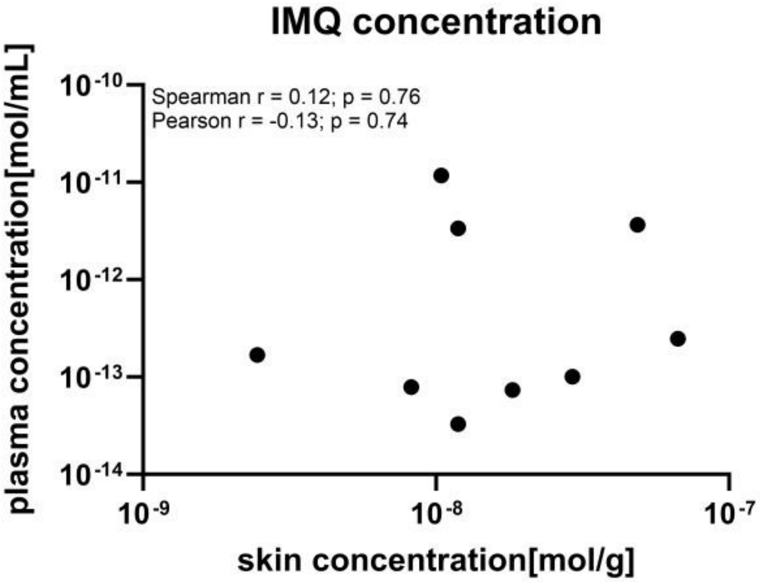


**Supplementary Figure 1. Lack of correlation between skin and plasma imiquimod (IMQ) concentrations.**

IMQ concentrations were measured in skin and plasma samples collected 24 h after the final topical application. Correlation analysis revealed no significant relationship between skin and plasma IMQ levels across individual animals ($n = 9$), as assessed by both Pearson ($r = -0.13$, $p = 0.74$) and Spearman ($r = 0.12$, $p = 0.76$) correlation tests. These data indicate that systemic exposure to IMQ does not directly reflect local skin concentrations under these experimental conditions.

| | Skin (IMQ compared to Control) | | | Plasma (IMQ compared to Control) | | |
|---|---|---|---|---|---|---|
| | Heatmap | Fold Change | p-value | Heatmap | Fold Change | p-value |
| **2-oleoyl glycerol** | ↑↑↑ | 2.918722538 | 8.88857E-08 | ↓ | -1.401011533 | 0.001461223 |
| **2-linoleoyl glycerol** | ↑↑↑ | 2.700263957 | 1.56769E-08 | ↓↓ | -1.818707545 | 1.41588E-08 |
| **2-arachidonoyl glycerol** | ↑↑↑↑ | 3.404962791 | 1.93288E-10 | ↓ | -1.468461411 | 2.82345E-05 |
| **palmitoyl ethanolamine** | ↑↑↑↑↑ | 23.32556788 | 2.22134E-07 | ↓ | -1.453647546 | 5.93721E-07 |
| **stearoyl ethanolamine** | ↑↑↑↑↑ | 20.05846132 | 3.55824E-09 | ↓ | -1.195161877 | 0.000167639 |
| **oleoyl ethanolamine** | ↑↑↑↑ | 5.655565563 | 4.42766E-06 | ↓ | -1.39016204 | 7.58233E-06 |
| **linoleoyl ethanolamine** | ↑↑↑↑↑ | 19.38007103 | 1.35635E-06 | ↓ | -1.324340528 | 0.001728771 |
| **arachidonoyl ethanolamine** | ↑↑↑↑↑ | 10.45827013 | 2.03298E-05 | ↓↓ | -1.5311523 | 1.93592E-07 |
| **docosahexaenoyl ethanolamine** | ↑↑ | 1.776074812 | 7.34455E-06 | ↓↓ | -1.578341014 | 2.40373E-08 |
| **eicosapentaneoyl ethanolamine** | BDL | BDL | BDL | | -1.048543689 | 0.783917369 |
| **palmitoyl alanine** | ↑↑ | 1.862004808 | 0.068946855 | ↓↓ | -1.571428571 | 0.007618624 |
| **stearoyl alanine** | ↑↑↑↑↑ | 11.20319672 | 0.000318984 | ↑↑↑↑ | 3.115909091 | 0.031390564 |
| **oleoyl alanine** | ↑↑↑↑ | 5.699316798 | 1.88268E-05 | ↓↓↓ | -2.008298755 | 1.229E-05 |
| **linoleoyl alanine** | | 1.02669866 | 0.876996917 | ↓↓ | -1.836477987 | 6.86882E-05 |
| **arachidonoyl alanine** | BAL | BAL | BAL | ↓↓ | -1.871323529 | 0.099428321 |
| **docosahexaenoyl alanine** | | 1.006670541 | 0.979548315 | ↓↓↓ | -2.072972973 | 0.005480451 |
| **oleic acid** | ↑↑↑↑↑ | 42.97549808 | 7.32103E-07 | ↓↓↓ | -2.0210317 | 4.0467E-09 |
| **linoleic acid** | ↑↑ | 1.730366781 | 0.000493444 | ↓↓ | -1.61832567 | 2.97639E-09 |
| **arachidonic acid** | ↑↑↑ | 2.477658049 | 7.68873E-06 | ↓ | -1.352745657 | 0.000166958 |
| **docosahexaenoic acid** | ↑↑↑ | 2.181067998 | 1.55098E-05 | ↓↓ | -1.65380185 | 1.2651E-06 |
| **stearidonic acid** | BDL | BDL | BDL | ↓↓↓ | -2.530577854 | 1.11469E-08 |
| **eicosapentaenoic acid** | ↑↑↑ | 2.001058668 | 6.92755E-05 | ↓↓ | -1.917244618 | 0.000126139 |
| **palmitoyl GABA** | | 2.104552969 | 0.115331659 | | -1.242352941 | 0.376347655 |
| **stearoyl GABA** | | 1.060720251 | 0.767191802 | BAL | BAL | BAL |
| **oleoyl GABA** | | 1.195687117 | 0.480133437 | BDL | BDL | BDL |
| **linoleoyl GABA** | BDL | BDL | BDL | BDL | BDL | BDL |
| **arachidonoyl GABA** | BDL | BDL | BDL | BDL | BDL | BDL |
| **docosahexaenoyl GABA** | BDL | BDL | BDL | BAL | BAL | BAL |
| **palmitoyl glycine** | | 1.177897378 | 0.321493922 | ↑↑↑↑ | 5.582037471 | 1.23246E-05 |
| **stearoyl glycine** | ↑↑↑↑ | 3.841637192 | 3.64626E-05 | ↑↑↑↑↑ | 41.64184242 | 1.47946E-09 |
| **oleoyl glycine** | ↑↑↑ | 2.625859249 | 0.00028128 | ↑ | 1.236320792 | 0.048934205 |
| **linoleoyl glycine** | | 1.356914538 | 0.178506994 | ↓ | -1.402708217 | 0.000638001 |
| **arachidonoyl glycine** | ↑ | 1.48881026 | 0.026508604 | | -1.177370031 | 0.318250991 |
| **docosahexaenoyl glycine** | | -1.382701061 | 0.172979369 | ↓↓ | -1.967032967 | 0.001076124 |
| **palmitoyl leucine** | ↑↑↑↑ | 4.105378389 | 1.53829E-05 | ↓ | -1.329889299 | 0.066979852 |
| **stearoyl leucine** | ↑↑↑↑↑ | 15.40379304 | 9.07514E-08 | ↑↑↑↑↑ | 338.7398374 | 2.3006E-12 |
| **oleoyl leucine** | ↑↑↑ | 2.71287599 | 0.002129914 | | -1.158461538 | 0.384432568 |
| **linoleoyl leucine** | ↓ | -1.395696034 | 0.031074483 | | -1.142719846 | 0.267435861 |
| **arachidonoyl leucine** | BAL | BAL | BAL | ↑ | 1.311688312 | 0.058281433 |
| **docosahexaenoyl leucine** | ↑↑↑↑ | 3.238663841 | 4.29725E-09 | | 1.103211009 | 0.231822106 |
| **palmitoyl methionine** | | 1.140611624 | 0.301763732 | ↓↓ | -1.907801418 | 0.019722929 |
| **stearoyl methionine** | BAL | BAL | BAL | ↑↑↑↑ | 9.196974281 | 7.94128E-08 |
| **oleoyl methionine** | BAL | BAL | BAL | ↓↓ | -1.530201342 | 0.000580176 |
| **linoleoyl methionine** | BAL | BAL | BAL | ↓↓ | -1.721048798 | 0.000694916 |
| **arachidonoyl methionine** | BDL | BDL | BDL | | -1.205830389 | 0.373048938 |
| **docosahexaenoyl methionine** | BDL | BDL | BDL | ↓↓ | -1.528783383 | 0.00723572 |
| **palmitoyl phenylalanine** | ↑↑↑↑ | 3.37664578 | 1.35239E-05 | ↓ | -1.402830189 | 0.010621165 |
| **stearoyl phenylalanine** | ↑↑↑↑↑ | 15.24184359 | 9.57813E-08 | ↑↑↑↑↑ | 45.46933962 | 2.22935E-12 |
| **oleoyl phenylalanine** | ↑↑↑ | 2.59231342 | 0.001568661 | | -1.079735683 | 0.570452661 |
| **linoleoyl phenylalanine** | | 1.329674778 | 0.192688617 | ↓ | -1.448497854 | 0.001418713 |
| **arachidonoyl phenylalanine** | | -1.131148844 | 0.591172355 | | -1.169590643 | 0.136561986 |
| **docosahexaenoyl phenylalanine** | ↑↑ | 1.521713628 | 0.005943708 | | -1.10475382 | 0.421199084 |

| | Skin (IMQ compared to Control) | | | Plasma (IMQ compared to Control) | | |
|---|---|---|---|---|---|---|
| | Heatmap | Fold Change | p-value | Heatmap | Fold Change | p-value |
| palmitoyl proline | ↓↓↓ | -2.16163954 | 0.00011556 | | 1.207172496 | 0.306534955 |
| stearoyl proline | ↑↑ | 1.620066721 | 0.006015296 | ↑↑↑ | 2.12244898 | 2.4586E-05 |
| oleoyl proline | | -1.034995075 | 0.779266602 | BDL | BDL | BDL |
| linoleoyl proline | BDL | BDL | BDL | BDL | BDL | BDL |
| arachidonoyl proline | BDL | BDL | BDL | BDL | BDL | BDL |
| docosahexaenoyl proline | BDL | BDL | BDL | BDL | BDL | BDL |
| palmitoyl serine | ↑↑↑ | 2.618235962 | 2.17255E-06 | | -1.036823105 | 0.799398377 |
| stearoyl serine | ↑↑↑↑ | 8.933033433 | 3.82483E-07 | | 1.17949111 | 0.500855208 |
| oleoyl serine | ↑↑ | 1.892191925 | 0.00017468 | ↓↓ | -1.811557789 | 0.052653018 |
| linoleoyl serine | ↑↑↑ | 2.350577031 | 0.001486713 | BDL | BDL | BDL |
| arachidonoyl serine | BDL | BDL | BDL | BDL | BDL | BDL |
| docosahexaenoyl serine | BDL | BDL | BDL | BDL | BDL | BDL |
| palmitoyl taurine | ↑↑↑↑ | 3.370718821 | 9.17072E-11 | ↑↑↑↑ | 7.986469557 | 6.30324E-11 |
| stearoyl taurine | ↑↑↑↑↑ | 15.66565859 | 7.56075E-09 | ↑↑↑↑↑ | 54.89169804 | 1.77373E-10 |
| oleoyl taurine | ↑↑↑↑ | 5.059702407 | 4.34239E-09 | ↑↑ | 1.973423487 | 1.48726E-08 |
| arachidonoyl taurine | BAL | BAL | BAL | | -1.025620687 | 0.592151352 |
| palmitoyl tryptophan | ↑↑ | 1.661947039 | 0.019189407 | | 1.272151899 | 0.191240329 |
| stearoyl tryptophan | BDL | BDL | BDL | | 1.100554236 | 0.421083873 |
| oleoyl tryptophan | ↑↑ | 1.910650689 | 0.081614871 | ↓↓ | -1.720765027 | 0.001059034 |
| linoleoyl tryptophan | BDL | BDL | BDL | ↑ | 1.44 | 0.040043379 |
| arachidonoyl tryptophan | BDL | BDL | BDL | BDL | BDL | BDL |
| docosahexaenoyl tryptophan | BDL | BDL | BDL | BDL | BDL | BDL |
| palmitoyl tyrosine | ↑↑↑↑ | 4.623998986 | 0.01421781 | ↓↓↓ | -2.01164295 | 0.006934002 |
| stearoyl tyrosine | BAL | BAL | BAL | ↑↑↑↑ | 3.134567901 | 0.001490956 |
| oleoyl tyrosine | BAL | BAL | BAL | ↓↓ | -1.613502935 | 0.002764378 |
| linoleoyl tyrosine | BAL | BAL | BAL | ↓↓↓ | -2.937904269 | 0.000239359 |
| arachidonoyl tyrosine | BDL | BDL | BDL | BAL | BAL | BAL |
| docosahexaenoyl tyrosine | BAL | BAL | BAL | ↓ | -1.442271881 | 0.008973345 |
| palmitoyl valine | ↑↑↑↑ | 5.83918333 | 0.000388727 | ↓↓ | -1.617231638 | 0.00606481 |
| stearoyl valine | ↑↑↑↑↑ | 18.74619627 | 2.14544E-05 | ↑↑↑↑↑ | 24.54903364 | 2.20899E-15 |
| oleoyl valine | ↑↑↑ | 2.087713391 | 0.001192982 | | -1.142050391 | 0.285596216 |
| linoleoyl valine | ↓↓ | -1.614242533 | 0.013715914 | ↓ | -1.355684211 | 0.002749785 |
| arachidonoyl valine | | 1.301019465 | 0.117367271 | | 1.527777778 | 0.1283573 |
| docosahexaenoyl valine | | 1.089066707 | 0.756081584 | ↓ | -1.4595186 | 0.029345516 |
| Prostaglandin E2 | | 1.204901394 | 0.350566154 | | -1.085527213 | 0.514969085 |
| Prostaglandin F2a | ↑ | 1.341299253 | 0.000164069 | | -1.11945926 | 0.18066132 |
| 12(S)-HETE | ↑↑↑↑ | 9.763188669 | 0.00048723 | ↓↓↓ | -2.15612272 | 0.060887167 |
| IMQ | | 103.4325086 | 0.100825165 | ↑↑↑↑↑ | 74.83786848 | 0.061937946 |
| Hydrocortisone | BDL | BDL | BDL | BDL | BDL | BDL |
| CA | ↓↓↓ | -2.973615301 | 0.043173326 | ↓↓↓↓↓ | -10.61813489 | 0.003001384 |
| DCA | ↑↑ | 1.589017527 | 0.037311076 | | -1.016370834 | 0.947402535 |
| GCA | | -1.026805806 | 0.933883198 | | -1.439444027 | 0.483300572 |
| TCA | ↑↑↑↑ | 5.176422075 | 2.02682E-05 | ↑↑↑↑ | 3.332942297 | 1.8869E-05 |
| TUDCA | BAL | BAL | BAL | ↑↑ | 1.732695931 | 0.016342458 |
| TDCA+TCDCA | ↑↑↑↑ | 3.46756868 | 0.004775529 | ↑↑↑↑ | 3.146677256 | 9.29682E-06 |
| GUDCA | | 1.560894105 | 0.141409568 | BDL | BDL | BDL |
| GDCA+GCDCA | | -1.782414586 | 0.292420714 | BDL | BDL | BDL |
| TLCA | BAL | BAL | BAL | | -1.227260813 | 0.560247685 |
| GLCA | BAL | BAL | BAL | BDL | BDL | BDL |
| Corticosterone | BDL | BDL | BDL | | 1.166012214 | 0.330324367 |

**Supplemental Figure 2: Data analysis of all endolipids scanned in skin and plasma.** The tables list the analytical data of the IMQ verses control comparisons. See Methods for description of Python data analysis for Fold change (denoted by arrows) and T-tests (colors denote significance levels and direction) to generate the Heatmap colors and arrow. Dark green represents significant increases after treatment ($p \leq 0.05$) while light green represents significance levels of $p \leq 0.1$-$0.05$. Dark orange represents significant decreases after treatment ($p \leq 0.05$) while light orange represents significance levels of $p \leq 0.1$-$0.05$. Fold change is indicated by the number of arrows, where 1 arrow corresponds to 1–1.49-fold difference, 2 arrows to a 1.5–1.99-fold difference, 3 arrows to a 2–2.99-fold difference, 4 arrows a 3–9.99-fold difference, and 5 arrows a difference of tenfold or more. BAL (below analytical limits) refers to endolipids that were measured in at least one sample but less than 4 making statistical analysis unreliable. BDL (below detectable limits) refers to endolipids that were not detected in any samples. Blank cells indicate that endolipids were present in at least 4 samples in each treatment group being analyzed and that no significant differences were present.
